# Molecular and biological characterization of an isolate of *Tomato mottle mosaic virus* (ToMMV) infecting tomato and other experimental hosts in a greenhouse in Valencia, Spain

**DOI:** 10.1101/063255

**Authors:** S. Ambrós, F. Martínez, P. Ivars, C. Hernández, F. de la Iglesia, S.F. Elena

## Abstract

Tomato is known to be a natural and experimental reservoir host for many plant viruses. In the last few years a new tobamovirus species, *Tomato mottle mosaic virus* (ToMMV), has been described infecting tomato and pepper plants in several countries worldwide. Upon observation of symptoms in tomato plants growing in a greenhouse in Valencia, Spain, we aimed to ascertain the etiology of the disease. Using standard molecular techniques, we first detected a positive sense single-stranded RNA virus as the probable causal agent. Next, we amplified, cloned and sequenced a ~3 kb fragment of its RNA genome which allowed us to identify the virus as a new ToMMV isolate. Through extensive assays on distinct plant species, we validated Koch’s postulates and investigated the host range of the ToMMV isolate. Several plant species were locally and/or systemically infected by the virus, some of which had not been previously reported as ToMMV hosts despite they are commonly used in research greenhouses. Finally, two reliable molecular diagnostic techniques were developed and used to assess the presence of ToMMV in different plants species. We discuss the possibility that, given the high sequence homology between ToMMV and *Tomato mosaic virus*, the former may have been mistakenly diagnosed as the latter by serological methods.

---

Vegetable crops, and particularly tomatoes, are one of the most important cultivated crops in Southern Europe. Spain is the main tomato producer in Europe, in terms of both total production (> 30%) and volume of fresh fruit exportation. The tomato production areas are centered in the Southeast of the country, with the region of Valencia being one of the main producers (Soler et al. 2010). Tomatoes are cultivated either in extensive greenhouses or in open fields, and several viral diseases cause important economic losses periodically (Soler et al. 2010). The frequent emergence or outbreaks of new viruses or of isolates of already known viruses reduces annual yield and quality of tomato production.

During the summer of 2015, virus-like symptoms consisting in epinasty, leaf distortion, foliar mottle, systemic crinkling, and fast necrosis were observed on 12 *Solanum lycopersicum* L. cv Moneymaker plants in a greenhouse near the city of Valencia (Fig. 1A and 1B). A virus was suspected as causal agent and total RNA extracts (RNAt) enriched in double-stranded RNAs (dsRNAs), were obtained by a standard RNA extraction protocol followed by a CF-11 chromatography (Ruiz-Ruiz et al. 2006). A dsRNA pattern, compatible with a viral infection induced by a positive single-stranded RNA virus (ssRNA), was observed in the extracts from symptomatic plants that was absent from the healthy ones (Fig. 1C). The dsRNA-rich extracts were reverse transcribed using a random hexamer primer (5’- GCCCCATCACTGTCTGCCCGNNNNNN-3’) (RACE technology) and *Moloney murine leukemia virus* retrotranscriptase (MoMLV RT; Promega, Madison WI, USA). cDNAs were then treated with Klenow DNA polymerase (Thermo Fisher Scientific Inc., Austin TX, USA) and amplified by a standard three-step PCR using the Phusion high-fidelity DNA polymerase (Thermo Fisher Scientific Inc., Austin TX, USA) and the primer 5’- GCCCCATCACTGTCTGCCCG-3’, as recommended by the manufacturer. Two different DNA band products of ~400 and ~650 bp were obtained, gel-purified (Zymoclean; Zymo Research, Irvine CA, USA), cloned and sequenced using standard molecular techniques.

**Fig. 1.**
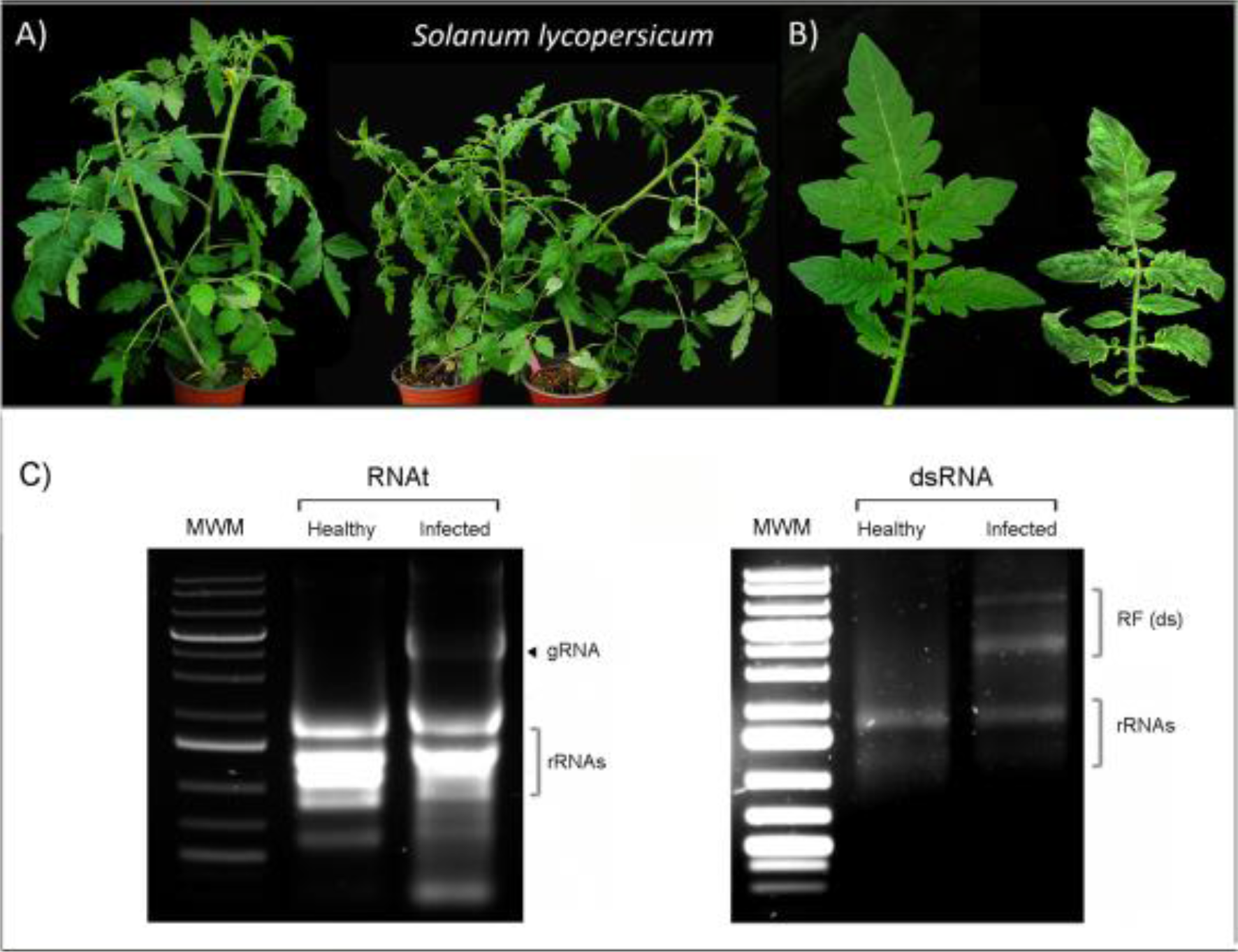
Symptoms of tomato (*S. lycopersicum* cv. Moneymaker) plants infected with the Spain/VLC-1 isolate of ToMMV and dsRNA characteristic pattern. A) Left: non-infected tomato plant. Right: infected tomato plant showing stunting, systemic mottling and apical necrosis. B) Detail of tomato leaves. Left: leaves from a non-infected plant. Right: leaves from an infected one showing leaf distortion, mottling, chlorotic flecks in younger leaves, and necrosis. C) RNA extracts from healthy (left lanes) or symptomatic tomato plants (right lanes) showing specific accumulation of ToMMV RNAs visualized by REAL safe staining (Durviz S.L., Valencia, Spain) in a native agarose gel. The positions corresponding to ToMMV genomic RNA (gRNA) in RNAt extracts (left panel) and the replicative dsRNA forms (RF) in dsRNA extracts (right panel) are indicated. rRNAs: plant ribosome RNAs. M: DNA molecular marker.

A BLAST search against the NCBI NR database revealed that one of the sequence fragments was 100% identical to a region of the RNA-dependent RNA-polymerase (RdRP) gene of the *Tomato mottle mosaic virus* (ToMMV) isolate Mexico/MX-5 (GenBank KF477193) in the genomic RNA positions 2766 to 3401. The second sequence showed 99% identity with same isolate, encompassing the 3’-terminal portion of the RdRP gene and the movement protein (MP) gene (positions 4767-5131). In order to fill the sequence gap between these two genomic fragments, five ToMMV-specific primers named RdRpF, RdRpR, 4800repR, MP, and MPR were designed based on the sequences obtained from Valencian isolate of ToMMV (Table 1). Standard RT-PCR and cloning protocols allowed us to obtain an assembled sequence with a final length of 3015 nt (deposited in GenBank with accession KU594507 and named Spain/VLC-1 isolate). This sequence has 19 nucleotide replacements with respect to the Mexico/MX- 5 isolate, all of them being transitions. Four of these transitions mapped into the MP gene, 3 of which being synonymous and 1 nonsynonymous. The partial fragment of the RdRP gene showed a total of 12 synonymous and 3 non-synonymous nucleotide substitutions, with two of the latter giving rise to non-conservative amino acid changes. To further explore the genetic similarity of ToMMV Spain/VLC-1 strain with other tobamoviruses, the homologous sequences from the three ToMMV isolates available at GenBank, Mexico/MX-5, USA/10-100 (KP202857) and USA/NY-13 (KT810183) were downloaded from the database. For comparative purposes, the homologous sequences from the three tobamoviruses showing the highest E-scores in a BLAST search against the NCBI NR database were also downloaded: *Tomato mosaic virus* (ToMV; KU321698), *Tomato brown rugose fruit virus* (TBRFV; KT383474) and *Tobacco mosaic virus* (TMV; EF392659). Sequences were aligned with CLUSTALX as implemented in MEGA5 (Tamura et al. 2011) and the nucleotide substitution model with the lowest *AIC* value, TN93 + G, was chosen for subsequent analyses. The average nucleotide divergence between Spain/VLC-1 and the three other ToMMV isolates is 0.007±0.002 (±1 SEM) for the genomic region analyzed. In contrast, the divergence from the other three tobamoviruses ranged between 0.220±0.012 for ToMV and 0.296±0.013 for TMV. A maximum likelihood tree, shown in Fig. 2, was constructed and its reliability evaluated with the bootstrap method (based on 1000 pseudoreplicates). All four ToMMV isolates form a highly significant monophyletic cluster clearly differentiated from the other three tobamoviruses. Within this cluster, the three ToMMV isolates from the New World are more closely related. In order to dispose of specific and reliable tools for rapid diagnostics of ToMMV, two molecular detection techniques based on an accurate RT-PCR test and on a hybridization protocol with digoxigenin (DIG)-labeled RNA probes were developed. In the case of the RT-PCR method, primers F98 and R99 (Table 1) were designed using the Spain/VLC-1 isolate as reference sequence. To validate its suitability, RNAt extracts from symptomatic and asymptomatic tomato plants growing in the greenhouse cabin where the virus was firstly detected, were used as test samples. Healthy controls obtained from tomato plantlets growing in a different cabin were also included. The RT-PCR protocol consisted in the following steps: (*i*) cDNA synthesis performed with the MoMLV RT following the manufacturer’s instructions, and, (*ii*) a three-step PCR amplification reaction performed with the Phusion DNA polymerase. Amplification cycling profile was 1 min at 98 °C for template denaturation, 5 cycles (98 °C 5 s, 68 °C 15 s, and 72 °C 15 s) using high *T_m_* annealing conditions followed by 35 additional cycles (98 °C 5 s, 65 °C 15 s, and 72 °C 15 s) for DNA amplification, and a final elongation step of 10 min at 72 °C. A 386 bp PCR product (Fig. 3A), specific indicator of ToMMV presence, was detected not only on the symptomatic tomato plants but also on a percentage of the asymptomatic plants from the original cabin (mainly juvenile plants). Since some suspicious yellowing symptoms were also observed in plantlets and juvenile plants of *Nicotiana benthamiana* Domin growing in different cabins of the same greenhouse, additional RT-PCR tests were also performed using RNAt from these plants (Fig. 3A left panel, lane 4). Plants showing these yellowing symptoms yielded the same PCR band (Fig. 3A left panel) and the specific presence of Spain/VLC-1 ToMMV isolate in this host was further confirmed by sequencing the amplicons.

**Table 1.**
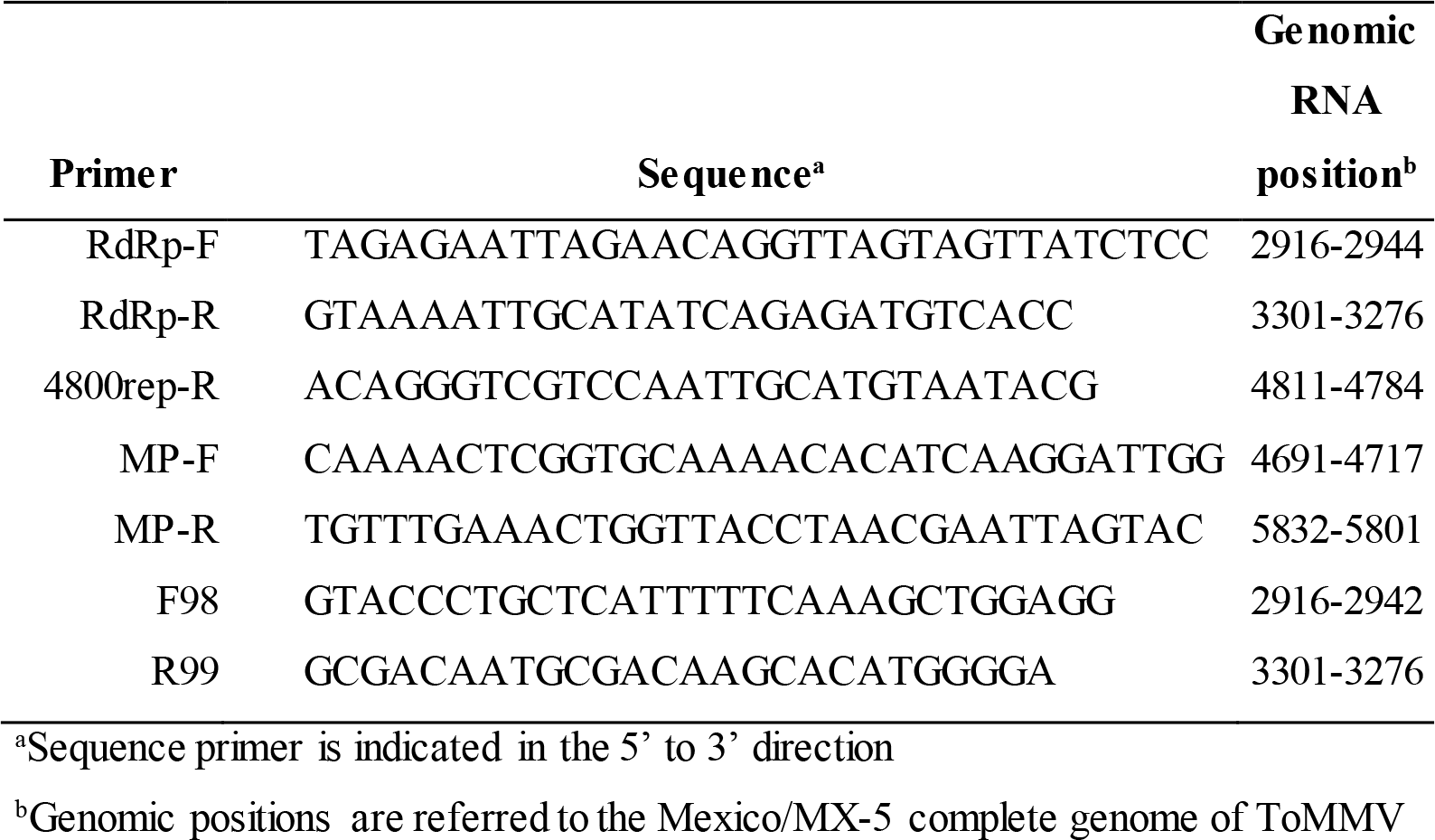
Primers used for ToMMV RT-PCR amplification reactions.

**Fig. 2.**
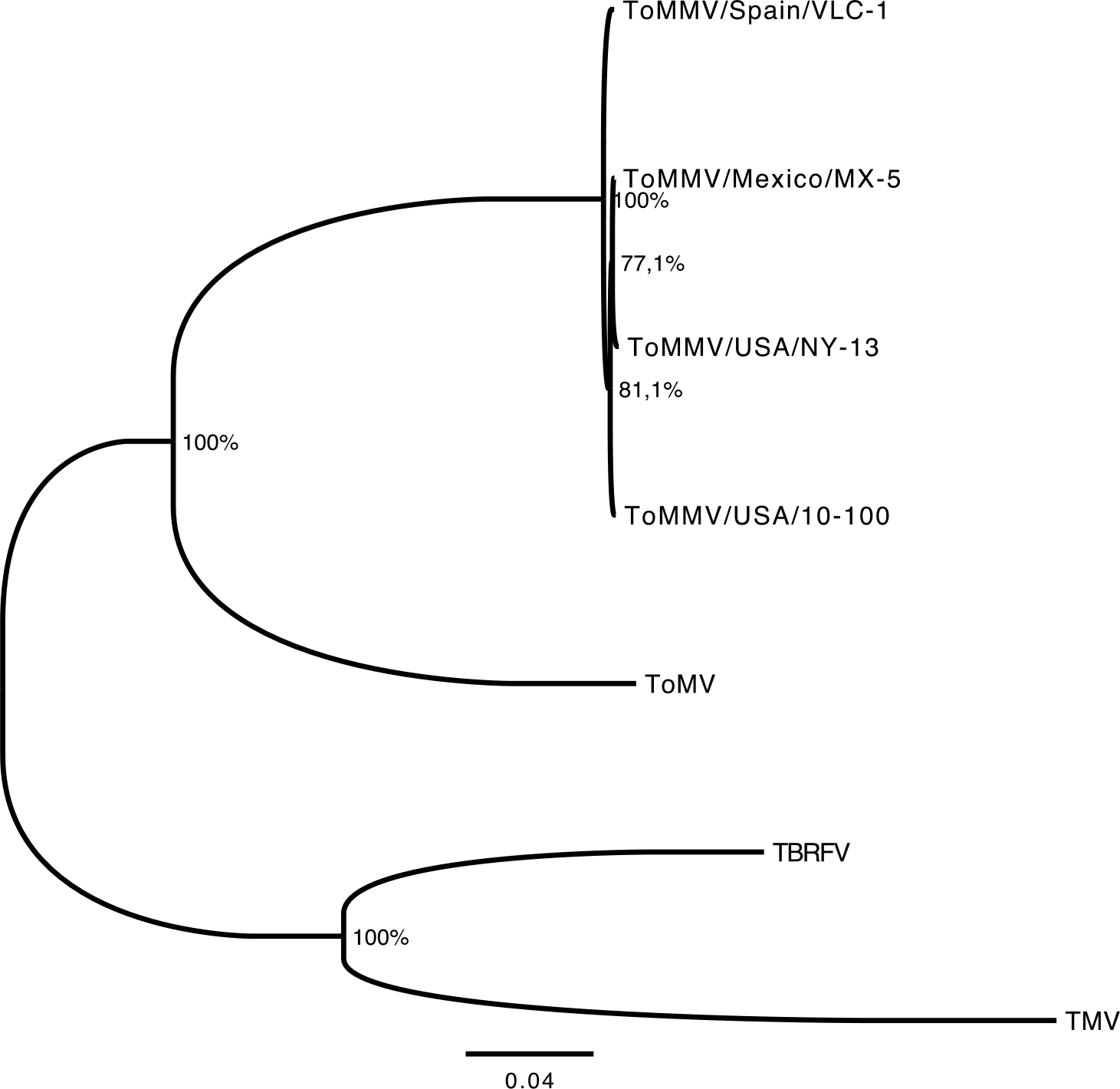
Maximum likelihood phylogenetic tree showing the relationship between ToMMV isolates and other members of the *Tobamovirus* genus. Only ToMMV isolates for which the sequence of the genomic region characterized in this study for Spain/VLC-1 isolate is known have been included in the analysis. All four ToMMV isolates form a highly significant monophyletic group based on the bootstrap support.

**Fig. 3.**
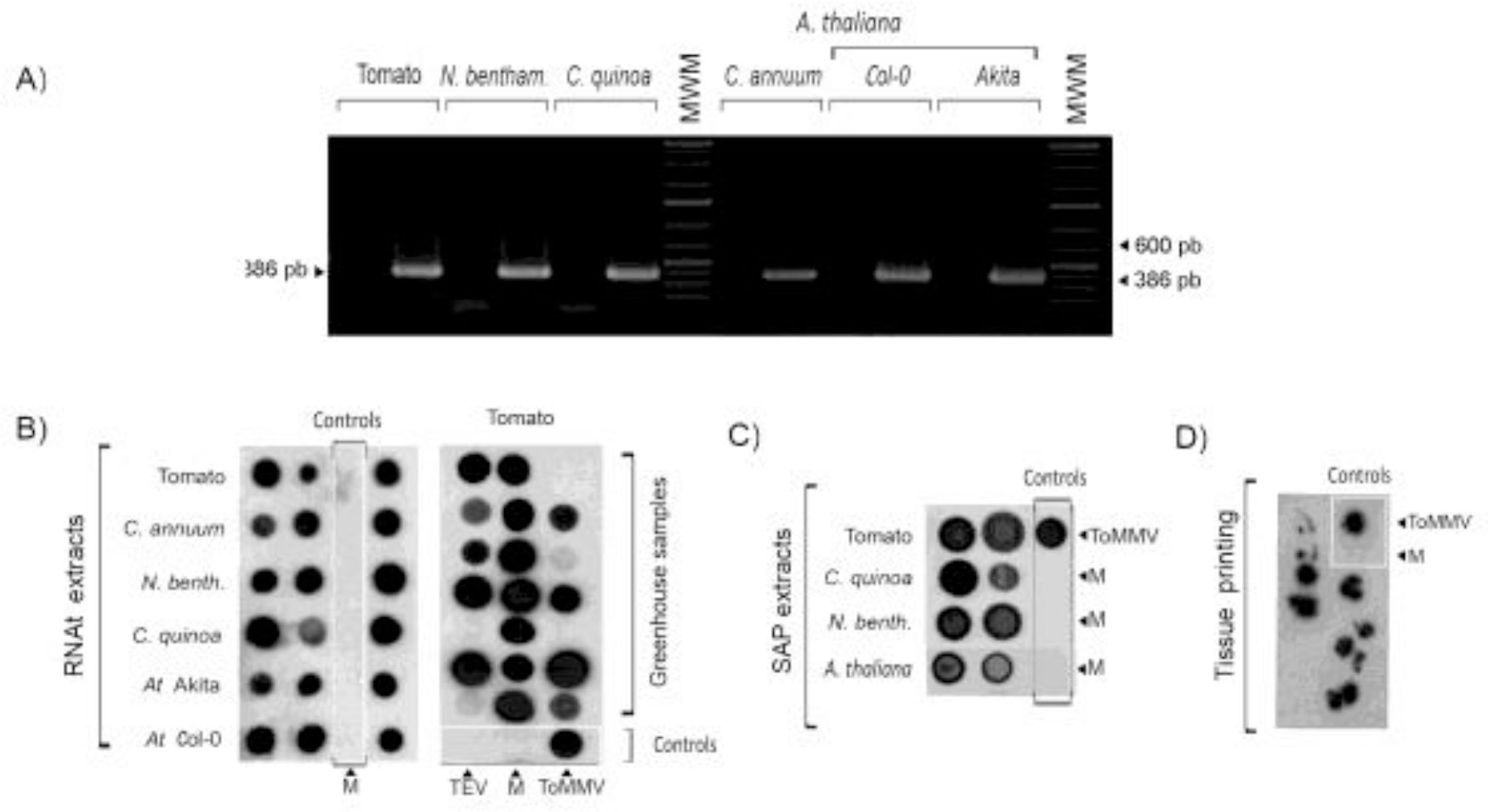
Detection of ToMMV with two molecular diagnostic methods. A) Left panel: RT-PCR detection using primers F98 (positions 2916 to 2944 in the gRNA) and R99 (positions 3301 to 3276). A specific band of 386 pb is shown. B and C) Detection of ToMMV by dot-blot molecular hybridization using RNAt (B) or sap extracts (C) and a DIG-RNA probe of negative polarity. D) Tissue printing hybridization of tomato greenhouse samples using the same approach. Control samples are delimited by a square line and an arrow. M: mock-inoculated plants, ToMMV: positive control, MWM: molecular weight marker, TEV: *Tobacco etch virus, N. bentham.:N. benthamiana,At:A. thaliana*.

For developing a ToMMV-specific northern hybridization assay, a specific ToMMV primer containing the T7 promoter sequence at the 5’ end of the R99 primer was designed and combined with the F98 primer for PCR amplification, using the specific DNA product of ToMMV obtained for the same region as template. An expected PCR product of 406 bp was obtained, that was purified and subjected to *in vitro* transcription with the T7 mMessage mMachine kit (Ambion Inc., Austin TX, USA). For the transcription reaction, the standard dNTP mix was replaced by the DIG-RNA labelling mix (Roche Diagnostics GmbH, Mannheim, Germany) to synthesize a ToMMV DIG-labeled RNA probe of negative polarity. The probe was used at a final concentration of 10 ng/mL in the hybridization solution (ULTRAHyb^©^; Ambion Inc., Austin TX, USA) and hybridizations were performed overnight at 68 °C following the manufacturer’s instructions. Samples consisting in tissue-prints, sap extracts or RNAt preparations from different plants were blotted onto positively-charged nylon membranes (Roche Diagnostics GmbH, Mannheim, Germany) and crosslinked. After chemioluminiscent detection, strong hybridization signals were observed in control positive samples and in most of the suspicious test samples (Fig. 3B and 3C), with no unspecific signals in negative controls nor cross-reaction signals in extracts from plants infected by other RNA viruses (Fig. 3B and 3C). These results confirmed the reliability of the hybridization approach and, moreover, corroborated the presence of ToMMV in different hosts.

Finally, to fulfill Koch’s postulates, we used sap extracts from the ToMMV-infected tomato source to inoculate new *S. lycopersicum* plants. Fifteen days later, the inoculated plants developed symptoms identical to those observed in the source plants. To go further with the biological characterization of the Spain/VLC-1 isolate, the same type of extracts was used to inoculate several plant species. We showed that ToMMV is sap-transmissible to a wide range of natural and experimental hosts as *Capsicum annuum* L., *N. benthamiana* L., *Nicotiana tabacum* L., *Arabidopsis thaliana* L., and the local-lesion host *Chenopodium quinoa* Willd. The following specific symptoms were observed: systemic mottling and necrosis in tomato (Fig. 1A), systemic crinkling and severe general chlorosis in *N. benthamiana* (Fig. 4B and 4C), fast yellowing and necrosis in pepper (Fig. 4D), and necrotic local lesions in *C. quinoa* (Fig. 4A). Several *A. thaliana* ecotypes were tested and although infection was confirmed by RT-PCR in all plants sampled (Fig. 3A, lanes 10 to 13), symptoms were only observed in Akita-0 (severe shrinking and leaf distortion; Fig. 4G) and Col-0 (some milder symptoms; Fig. 4E and 4F). This is the first description of ToMMV infecting and inducing symptoms in *A. thaliana*, a plant species extensively grown in research greenhouses as the prototypical experimental model system in plant biology.

**Fig. 4.**
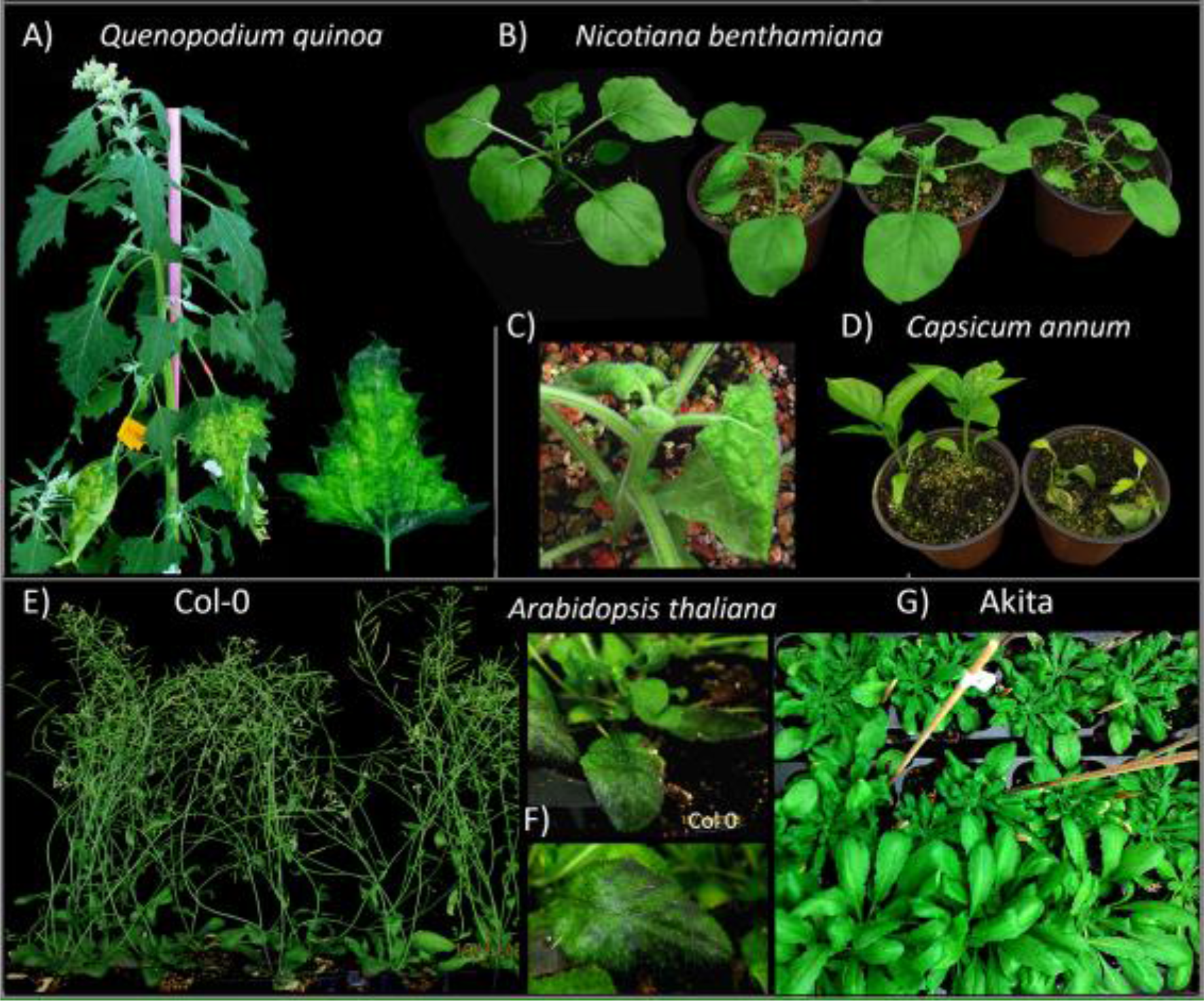
Biological characterization of ToMMV Spain/VLC-1 isolate in different host species. A) Local-lesion host *C. quinoa*. Detail of a leaf showing severe local lesions. B) Left: noninfected experimental host *N. benthamiana*. Right: infected *N. benthamiana* plants showing severe stunting, systemic crinkling and leaf distortion with apical necrosis. C) Detail of an apical shoot from an infected *N. benthamiana* plant. D) Left: non-infected *C. annuum* cv. Largo de Reus plant. Right: infected *C. annuum* plant showing severe apical shoot yellowing and necrosis. E to G) Experimental host *A. thaliana* accessions infected with ToMMV. E and F) mild stunting and leaf curling with dark brown flecks observed in accession Col-0 (non-infected plant shown at right). Severe stunting, systemic crinkling and leaf malformation in accession Akita-0 (infected plants at the top showing severe epinasty, non-infected plants at the bottom).

The genus *Tobamovirus* is worldwide distributed and actually includes more than 30 members that infect several host species. Among them, TMV and ToMV are the most common viruses infecting solanaceous species of agronomic importance, causing important damages in infected plants. Both are very efficiently transmitted mechanically and by seeds, contributing to their fast spread (Lewandowski and Dawson 1998; Hadas et al. 2004). ToMMV was first reported by Li et al. (2013) as a novel *Tobamovirus* species infecting tomatoes from a greenhouse in Mexico, and it has been recently described in China (Li et al. 2014), USA (Webster et al. 2014; Filmer et al. 2015; Padmanabhan et al. 2015), Brasil (Moreira et al. 2003), and Israel (Turina et al. 2016). Additionally, Pirovano et al. (2015) revealed its presence in an ancient variety of *Cicer arietinum* L. in Italy by using a metagenomics approach. Here, we are reporting ToMMV infection in Spain. From its initial description, ToMMV has become a major concern for tomato industry, as it was ToMV. In fact, ToMV is greatly widespread among tomato production areas, and often causes epidemics in tomato and pepper cultivars. In Valencia, ToMV and *Potato virus Y* (PVY) were described as the two most prevalent viruses in tomatoes (Soler et al. 2010). Given the high amino acid sequence similarity between ToMV and ToMMV, it will not be surprising that many of the infections that are being diagnosed as ToMV using serological methods would actually be ToMMV, as was already suggested by Turina et al. (2016). Therefore, the real incidence and epidemiology of this emerging virus needs to be undertaken.

## Acknowledgements

This work was supported by grants BFU2015-70261-P and BFU2015-65037-P (to C.H. and S.F.E., respectively) from Spain Ministry of Economy and Competitiveness.

